# A novel two-generation approach for understanding the population dynamics of gregarious parasitoids

**DOI:** 10.1101/2021.02.22.432341

**Authors:** Alena Samková, Jan Raška, Jiří Hadrava, Jiří Skuhrovec

## Abstract

Parasitoids, as important natural enemies, help maintain balance in natural ecosystems. Their population dynamics is generally predicted from the number of individuals. Here, using gregarious parasitoids as models, we show that this traditional approach omits one important parameter: the mother’s manipulation of offspring fertility due to the clutch size–body size– fertility correlation. As a result of this correlation, when females deliberately adjust the number of offspring laid in a host, they determine not only the number but also the body sizes and reproductive potentials of those offspring. For the first time in the model species *Anaphes flavipes*, we determined the parasitoid’s offspring fertility from clutch size. Using this, we experimentally clarified the advantage of specific clutch size combinations and we show that identical fertility in the F1 generation can lead to distinctly different fertility values in the F2 generation. Even with the same number of hosts, lower fertility in the F1 generation can cause higher fertility in the F2 generation. Based on these results, we propose a novel two-generation approach which includes the clutch size–body size–fertility correlation. Our novel approach provides a new perspective for determining the individual fitness levels of gregarious parasitoids with new options for the modelling of parasitoid population dynamics.

## INTRODUCTION

The diversity of interspecific interactions between organisms is a remarkable natural phenomenon. One of these relationships is the parasitoid-host interaction, common in insects^1^. The females of parasitoids lay their eggs into various developmental stages of arthropods; their larvae develop in the host, feeding on its tissues and killing it before completing their own development^2^. Due to this strategy, parasitoids play an important role as natural enemies in maintaining the biodiversity and balance in natural ecosystems^3,4^. They occur in high numbers, both in terms of species diversity and absolute numbers of individuals^5^.

From the ecological point of view, all parasitoids can be divided into two groups, solitary and gregarious, according to their reproductive strategy^6^. The solitary strategy is ancestral and the majority of known species of parasitoid wasps develop solitarily^7^. Their females lay one or more eggs into a single host, their predatory and more mobile larvae compete until only one survives^8,9^. The gregarious strategy is derived and has evolved numerous times (independently at least 43 times in 26 different families of Hymenoptera)^10,11^. In contrast to the solitary larvae, those of gregarious parasitoids are tolerant, therefore multiple individuals may develop in a single host^8,9^. The tolerance model of Godfray^10^ assumes that when sharing the same host, tolerant gregarious larvae will always be killed by a predatory solitary larvae (examined by Laing & Corrigan^13^). It is therefore assumed that the gregarious strategy must provide their holders with advantages that compensate for this handicap.

One of the main benefits of the gregarious strategy is the efficient use of the host in the form of multiple individuals developing in a single host. However, for many gregarious parasitoids, a higher clutch size reduces the body size of the offspring^e.g.11,14^ and their future fertility (i.e., the clutch size – body size – fertility correlation)^15,16^. In optimal conditions, the females distribute their offspring among as many hosts as possible. However, with a scarcity of hosts, the mothers face a trade-off between the body size and number of their offspring: by laying a different clutch size they choose either more, smaller offspring with a lower fertility, or fewer, larger offspring with a higher fertility^16^.

Some clutch sizes or their combination are more advantageous than others^17^ and therefore favoured by natural selection. Over time these advantageous clutch sizes should stabilize to maximize individual fitness^17,18^; this would indicate the existence of *a stable reproductive strategy*. However, more recent studies suggest the ability of the female to assess the variance in external conditions and change their reproductive strategy (including clutch size) accordingly^19,20,21^ – *the variable reproductive strategy*.

It is obvious that predicting the parasitoid’s population dynamics from a traditionally approach, i.e. determination from the number of individuals and then from maternal fertility^22,23^, is suitable for solitary parasitoids, where only one offspring completes larval development in one host and its body size and future fertility are directly proportional^e.g.24^. However, for predicting the population dynamics of gregarious parasitoids with the variable reproductive strategy and varying fertility, this traditionally approach may provide inaccurate results.

Here, in the model species for gregarious parasitoids, *Anaphes flavipes* (Förster, 1841), for the first time to our knowledge, we experimentally showed the efficiency of host utilization in terms of acquired fertility of offspring from different clutch sizes. Using this, we experimentally demonstrated the advantage of specific clutch size combinations and examined whether females under optimal conditions distribute their offspring randomly or prefer advantageous combinations.

In the second part in this study, we proposed a novel, two-generation approach for determining the individual fitness levels of gregarious parasitoids. We show that using this approach we are able to obtain a more accurate view of the population dynamics of gregarious parasitoids, compared to the traditional approach of evaluating the maternal fertility only.

## MATERIALS AND METHODS

### Parasitoids

*Anaphes flavipes* were reared from host eggs (*Oulema* spp.) collected in cereal fields in Prague (50.136°N, 14.363°E) from the end of April until the end of June in 2012, 2019 and 2020. The parasitised host eggs were stored in Petri dishes with moistened filter papers until adult wasps emerged. These “wild” wasps were used as an initial population from which the next generations of parasitoids were reared in an environmental chamber at 22±2°C with 40– 60% relative humidity and continuous illumination. Subsequent generations of females and males were used for experiments. Mated females (not older than 24 hours post-emergence) were placed in Petri dishes with host eggs. The females were not fed before the start of the experiment or during the experiment, and they had free access to water (modified from Samková et al.^16,25^).

### Host species

The non-native host species *Gastrophysa viridula* (DeGeer, 1775) and native hosts of the *Oulema* species complex (including two very ecologically close species, *O. duftschmidi* (Redtenbacher, 1874) and *O. melanopus* (Linnaeus, 1758)) were used; these species were used identically as in our previous studies^e.g.16,25^ and in other studies^e.g.26,27^ because they are determined only on the basis of genital preparation^28^. In the current study, the host culture was established from adults collected in Prague (50.136°N, 14.363°E) and in Police n/Met (GPS: 50.527°N, 16.245°E). The adults were kept in plastic boxes with moistened filter papers, were fed, and had unlimited access to water. They were allowed to lay their eggs on leaves at 22±2°C, a relative humidity of 40–60% and a 16:8-h L:D cycle. We used host eggs no older than 24 hours (modified from Samková et al.^16,25^).

#### Laboratory experiments

All laboratory experiments were performed in Petri dishes (8.5 cm) in a thermal cabinet at 22±2°C and 40–60% relative humidity under a 16:8-h (L:D) photoperiod. Individual parasitised host eggs were moved into 1.5-ml plastic tubes on the 9^th^ or 10^th^ day after parasitisation and stored at the same temperature in a thermal cabinet. The number and sex ratio of wasps that emerged from each parasitised host egg were measured. The body size of the parasitoids was measured using temporary microslides according to established methodology^16^ (modified from Samková et al.^16,25^).

##### Exp. 1) Determining fertility in the F2 generation

Each female (n=81) had 12 host eggs available for parasitisation for 8 hours. We measured the body sizes of the females and their fertility values to determine the relationship between body size and fertility. After this, we measured the clutch size and body sizes of the offspring (n=441) to determine fertility from the body size of *A. flavipes*. Data from these experiments were used for studies (Samková et al. 2019ab) focused on the body size, fertility and factors influencing the clutch size of *A. flavipes* female parasitoids (Suppl. Mat. 1).

##### Exp. 2) Determining advantageous combinations of clutch sizes

Each female (n=19) had 12 host eggs available for parasitisation for 8 hours. We determined the fertility values of the offspring in the F2 generation of each female. While maintaining the number of parasitised host eggs and the number of offspring (without male fertility), we determined the most advantageous (highest fertility values in the F2 generation) and least advantageous (lowest fertility values in the F2 generation) combinations for each female and compared them with reality (Suppl. Mat. 2).

##### Exp. 3) Effect of non-native hosts on clutch size

Each female had host eggs available for parasitisation for 8 hours in three groups: 1) 12 native hosts (n=19); 2) 6 native hosts (n=11); and 3) 6 non-native and 6 native hosts (n=15) (Suppl. Mat. 3).

#### Statistical data processing

We estimated the effect of the F0 female reproductive strategy on F1-generation fertility in a two-step analysis. First, we modelled linear regression between *A. flavipes* female wing length and fertility (both with normal distributions) based on measurements of wasps used in a previous study^16^. The parameters of the regression curve were calculated in Statistica 7 (StatSoft, Inc., 2004). In the second step, we calculated the hypothetical fertility of each female wasp used in the current study by applying regression parameters to their known wing lengths. The total wasp fertility per host was calculated by multiplying the sum of the hypothetical fertilities of the female offspring of each host by the mean rate of female offspring for each level of the total number of siblings per host (levels: 1, 2, 3, 4). Since the ultimate aim of this model was to estimate the population dynamics of the wasps, male fertility was considered to be zero.

To test the hypothesis that F0 females do not lay eggs into hosts randomly but instead select egg distributions that provide higher total fertility to the F1 generation, we compared the median F1 fertility of all possible combinations for each female with the one actually selected by the female by means of a paired t-test. The test was performed in Statistica 7 (Statsoft, Inc., 2004).

Analyses testing the effects of the number of native/non-native hosts on reproduction of *A. flavipes* reproduction were performed in R 4.0.3^29^. For the number of offspring per dish, we used linear models (LM), the dependent variable was square-root transformed if necessary (labelled as “LM-sqrrt”). The clutch size was analysed by means of mixed effect linear models (LMM) or generalized mixed-effect linear models for Poisson distribution (GLMM-p), depending on distribution of analysed subset, with an ID specific for each dish as a random factor. Mixed-effect models were built in R package lme4^30^ and analysed by their comparison with corresponding null models (including the random factor only) by means of ANOVA. All models included two-level subset of Group (levels: 6 native hosts; 12 native hosts; mixed hosts – 6 native and 6 non-native ones) as a single fixed factor. 95% confidence intervals displayed in Fig. 3 were calculated in R 4.0.3^29^ for normal data distribution and R package DescTools for Poisson distribution (Group levels “12 native hosts” and “mixed hosts”).

**Fig. 3.**
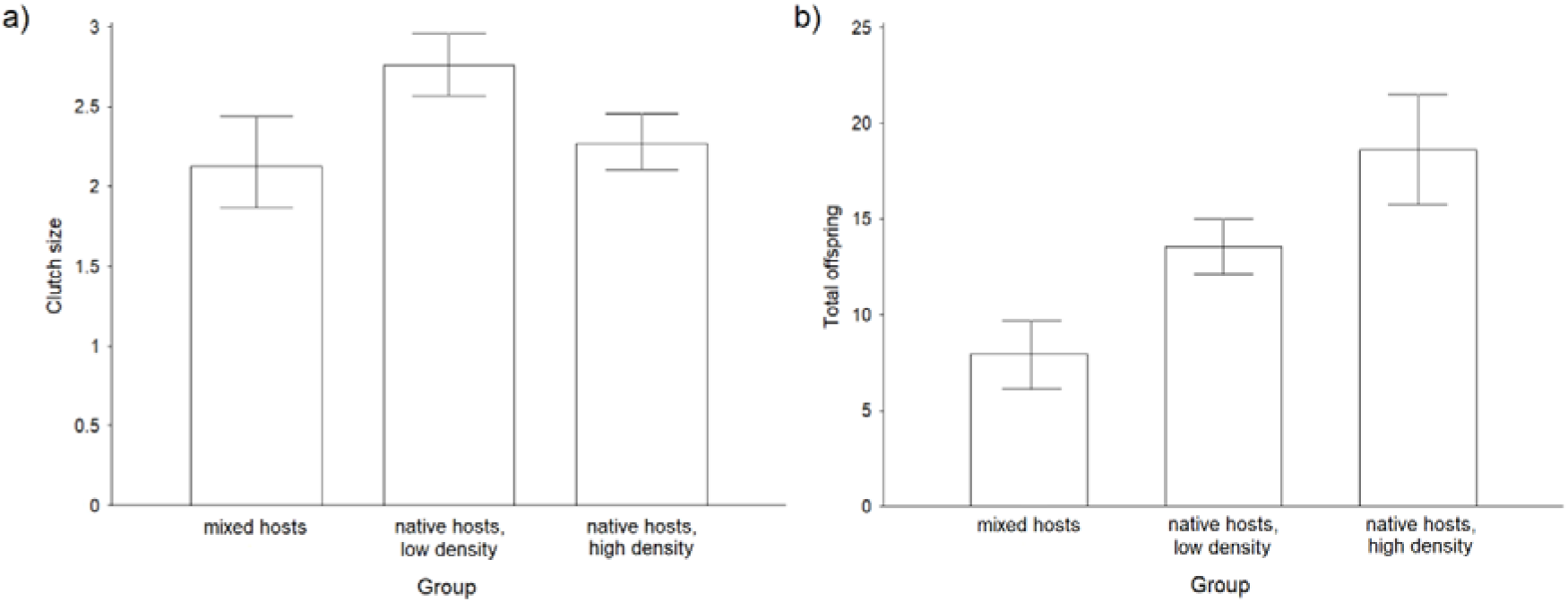
(A) Clutch size (i.e. offspring per parasitised host egg) and (B) total number of offspring per female *A. flavipes* under different conditions: mixed hosts (6 native hosts eggs and 6 non-native hosts eggs), native host, low host density (6 native hosts eggs); native host, high population density (12 non-native host eggs).

## RESULTS

### Exp. 1) Determining fertility in the F2 generation

The fertility of the gregarious parasitoid *Anaphes flavipes* among different clutch sizes is shown in Table 1 (Suppl. Mat. 1).

**Table 1.** The fertility of the gregarious parasitoid *A. flavipes* among different clutch sizes (mean ± 95% confidence interval). The column “c)” shows that the fertility of females in the F2 generation can be reduced by almost half, due to the different distribution of 12 offspring into clutch size.

| Clutch size | a) Fertility of one offspring | b) Fertility of all offspring per host | c) Fertility of 12 offspring |
| --- | --- | --- | --- |
| 1 | 19.3 ± 1.3 | 19.3 | 232 (100%) |
| 2 | 13.8 ± 0.3 | 27.6 | 166 (72%) |
| 3 | 11.2 ± 0.4 | 33.6 | 134 (58%) |
| 4 | 10 ± 1.5 | 40 | 120 (52%) |

Using this, we determined the host utilisation by gregarious parasitoids compared to solitary parasitoids from the novel two-generation approach (Fig. 1).

**Fig. 1.**
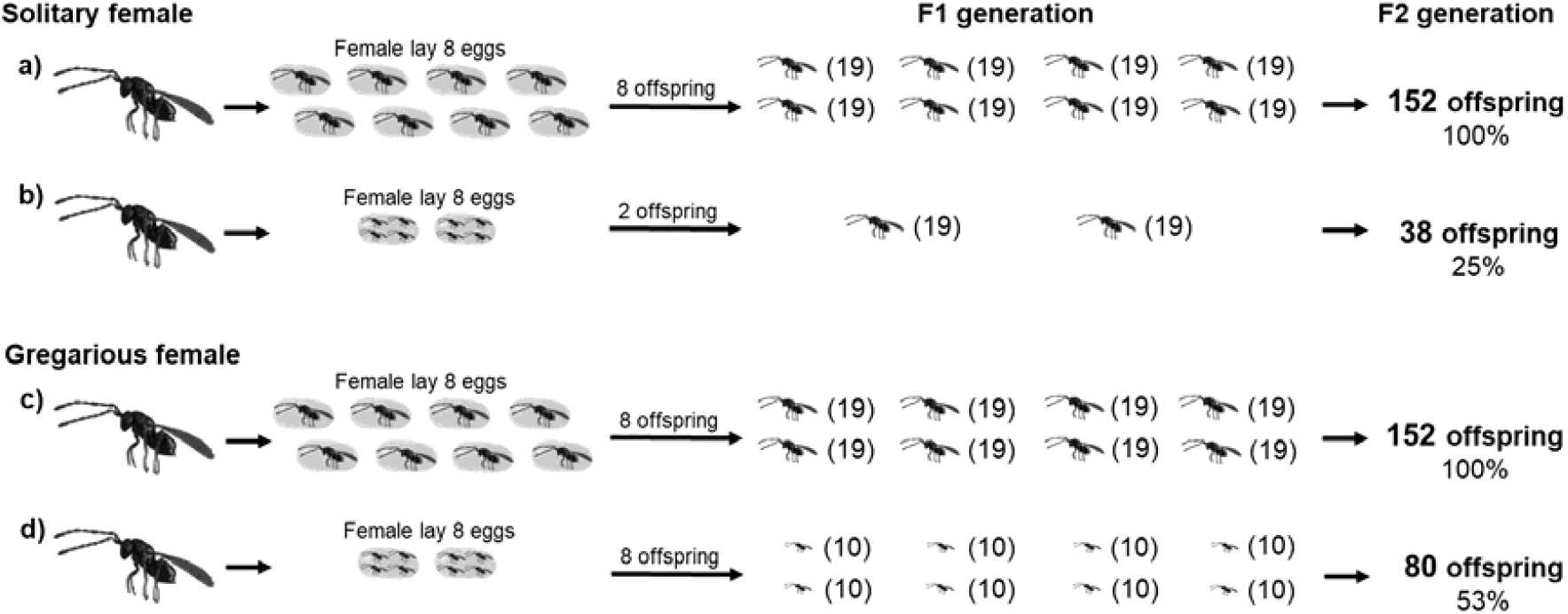
The different fertilities obtained in the F2 generation for solitary and gregarious parasitoids with different numbers of host eggs (to simplify the model, the fertility values (shown in parentheses) are rounded, and male offspring are not shown; these simplifications are explained in detail in the methods section *Exp. 1*).

### Exp. 2) Determining the advantageous combinations of clutch sizes

We experimentally showed that when parasitising a particular number of hosts, females did not distribute their offspring randomly, but chose combinations that increased the total fertility of the F1 generation (paired t-test, t = 8.932, p < 0.001) (Suppl. Mat. 2). Figure 2 shows that using advantageous combinations of clutch size (Fig. 2b), the females can obtain high fertility values in the F2 generation from same fertility in the F1 generation (Fig. 2a,b) and that, even with the same number of hosts, lower fertility in the F1 generation can cause higher fertility in the F2 generation (Fig. 2c).

**Fig. 2.**
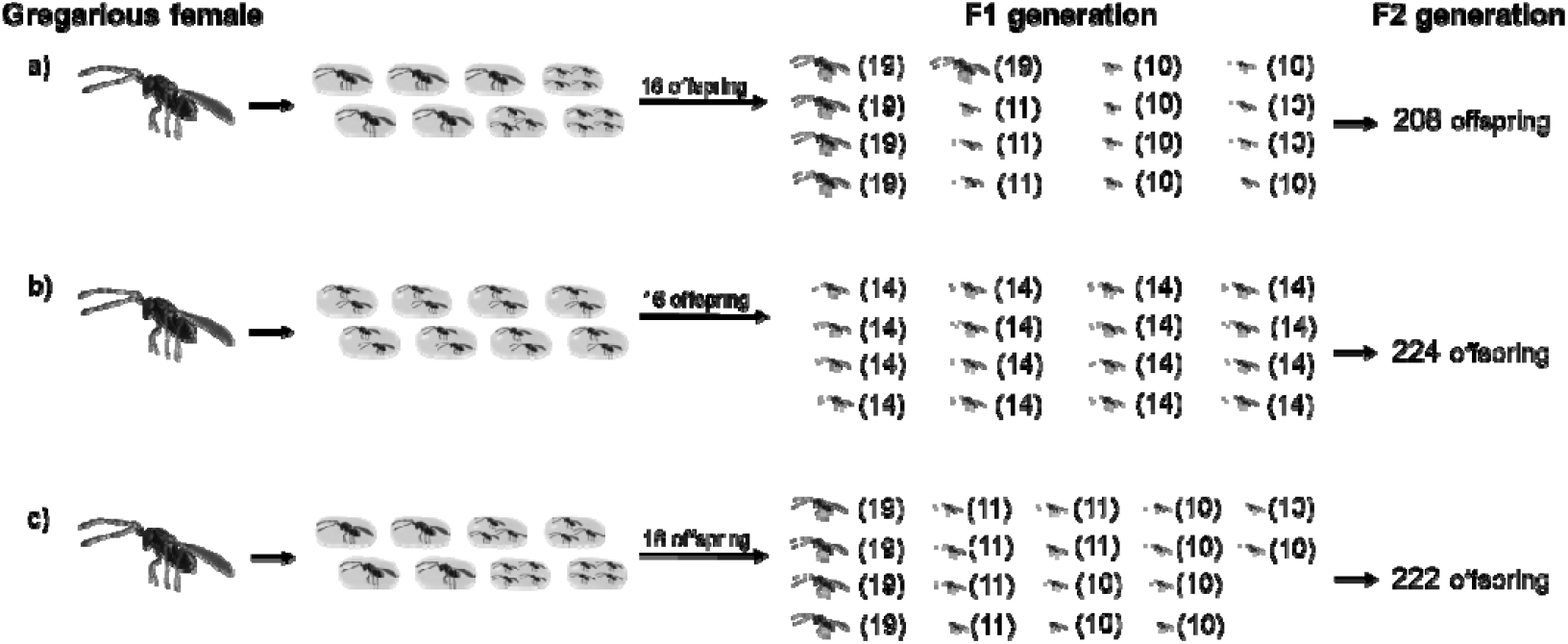
The different fertility values obtained in the F2 generation of females with the same number of host eggs and the same (2a, b) or different fertility values in the F1 generation (2c) (to simplify the model, the fertility values (shown in parentheses) are rounded; male offspring are not shown, explained in detail in the methods section *Exp. 2*).

### Exp. 3) Effect of non-native hosts on clutch size

The clutch sizes of *A. flavipes* females with 12 host eggs available for parasitisation were lower than those of females with 6 available host eggs due to the distribution of offspring among multiple hosts (LMM, χ^2^ = 6.824, p = 0.009) and the fertility between these two experimental groups was not statistically significant (LM-sqrrt, F_(1,29)_ = 5.754, p = 0.023; Fig 3). Females with 6 native hosts and 6 non-native hosts in which the offspring were unable to complete larval development had lower clutch sizes than females with 6 native hosts (LMM, χ^2^ = 12.193, p < 0.001) and no significant difference in clutch size was observed compared to females with 12 native hosts (GLMM-p, χ^2^ = 0.516, p = 0.473; Fig. 3). The fertility in the group where females had 6 native and 6 non-native host for parasitisation were lower compared to both the group of females with 6 native hosts (LM, F_(1,24)_ = 20.356, p < 0.001) and that of females with 12 native host (LM, F_(1,33)_ = 32.583, p < 0.001; Fig. 3) (Suppl. Mat. 3).

## DISCUSION

The evolution of parasitoids clutch size is one of the most enduring topics in behavioural ecology and life history theory^11^. For gregarious parasitoids, some combinations of the clutch size confer more advantages than others^17,31^, and are therefore favoured by natural selection to maximize individual fitness^18^.

In this study, we have clarified the advantage of clutch size combinations in the form of a higher fertility obtained in the F2 generation. Although a parasitoid’s fertility generally depends on the host’s population density and the number of eggs that a female has available^2^, we show that even with the same number of hosts and the same number of offspring, females of gregarious parasitoids can deliberately manipulate fertility in the F2 generation by choosing suitable clutch size combinations in the F1 generation (Fig. 2a, b). Moreover, with the same number of hosts, lower fertility in the F1 generation can cause a higher fertility in the F2 generation (Fig. 2b, c). Interestingly, we have shown that the wasps did not distribute their offspring randomly among a given number of hosts, but rather chose combinations that could be described as preferred^16,26^. We experimentally showed that using these combinations they increased total fertility in F2 generation.

Based on our results, we propose the novel two-generation approach, which takes into account the clutch size – body size – fertility correlation. Using this approach, we can obtain a more accurate view of the population dynamics of gregarious parasitoids compared to the traditional approach of maternal fertility^22,23^. For an illustration, we present a hypothetical scenario in which the same number of equally large *A. flavipes* parasitoids are released for biological control at two sites with different numbers of hosts (low and high). The population densities of the parasitoids, measured by abundance in the F1 generation (12 – 14 days after parasitoid release^26^), would be approximately the same at both sites, and differences would only be observed in the body size of the F1 offspring^25^. Using the two-generation approach, we know that the size of the parasitoid population in the F2 generation (24 – 28 days after parasitoid release^26^) would be notably lower at the site with a low host population density due to the reduced body sizes of offspring in the F1 generation. Using offspring fertility determined from the clutch size, one can predict population dynamics in these localities (Fig. 1c, d).

We are aware that determining the parasitoid fertility from clutch size requires extensive measurements difficult to apply in the field. As an alternative, we propose obtaining knowledge of key factors, in the case of *A. flavipes* varying host population densities, intraguild predation^25^ or superparasitism^32^, whereby the parasitoid female “intentionally” changes the clutch size. Understanding these factors in a specific environment, we can expect a decrease or increase in clutch sizes and offspring fertilities, i.e. specifically predict their population dynamics.

Interestingly, we have also found that clutch size can be affected “unintentionally” by the presence of non-native hosts in which the offspring of parasitoids *A. flavipes* are unable to complete larval development. Using the traditional one-generation approach, we would expect a reduced fertility by almost half in locations with a non-native host (Fig. 4). However, from a two-generation approach, we still expect reduced fertility on this locality, but only to a limited extent, because females, by non-random offspring distribution in both the non-native and native host, reduce their clutch size in the native host and thus increase the individual offspring fertility (Fig. 4c compared to 4b). If we apply the two-generation approach, measuring certain clutch sizes, we find that in these circumstances the fertility of individuals in the F1 generation in the presence of a non-native host decreases by 50% (Fig. 4c), but among individuals in the F2 generation, fertility decreases by only 27% (Fig. 4c).

**Fig. 4.**
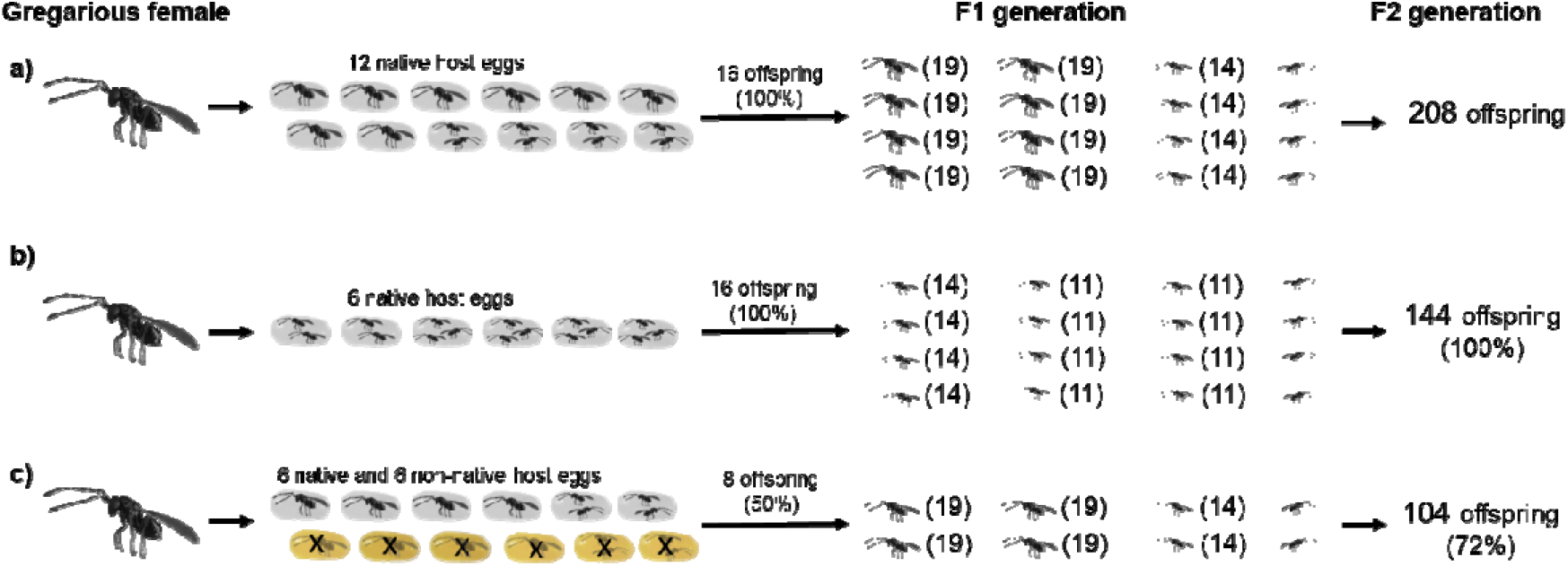
The different fertility values obtained for the F2 generations of three groups of females: a) females with 12 available native host eggs; b) females with 6 available native host eggs; and c) females with 6 available native host eggs and 6 available non-native host eggs (the offspring sex ratio of *A. flavipes* is 3:1 (male:female)^26^; females are shown as 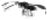 (left-facing) and males as 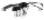 (right-facing)).

## CONCLUSION

The reproductive strategy of gregarious parasitoids is a particularly interesting biological puzzle, as it represents a fertility-dependent, flexible life history strategy. This strategy brings benefits to its users in the form of the plasticity of clutch size combinations, by which the parasitoids can “intentionally” and “unintentionally” manipulate the fertility of offspring in the future generation. Our novel two-generation approach, which takes these clutch size – body size – fertility correlations into consideration, allows more accurate results for predicting parasitoids’ population dynamics. We suggest that determining fertility from clutch size is useful for important species of gregarious parasitoids, and propose the two-generation approach as a model that may ensure more effective uses of parasitoids in biological plant protection or more successfully predict ecosystem stability.

## Supporting information

Suppl. mat. 1

Suppl. mat. 2

Suppl. mat. 3

## Acknowledgments

We would like to thank our colleagues who provided us with technical equipment: Jana Mazáková, Jana Wenzlová, Marie Maňasová, Pavel Ryšánek and Miroslav Zouhar. Finally, we thank Marek Romášek and AJE company for proofreadings the manuscript. This work has been supported by the Ministry of Agriculture of the Czech Republic, institutional support MZe-RO0418 and from programme NAZV No. QK1910281 (MZe ČR) (both to JS).

